# Advancing Oceanic Studies with HyperOCR Sensors and Non-Negative Matrix Factorization: A Cost-Effective, Data-Driven Approach for Analyzing Light in Marine Water Column

**DOI:** 10.1101/2024.05.25.595916

**Authors:** Mateo Sokač, Staša Puškarić

## Abstract

Understanding the intricate dynamics of ocean biogeochemistry is crucial for deciphering its role in climate change. Our study addresses this challenge by integrating advanced computational techniques and innovative sensor technology to enhance remote sensing capabilities. Drawing on recent insights into the vast carbon reservoirs within the ocean, particularly within the dissolved organic matter (DOM) pool, we highlight the pressing need for comprehensive spatial and temporal understanding facilitated by a combination of satellite and in situ data. However, existing remote sensing methods face limitations in capturing subsurface processes, hindering our ability to grasp carbon fluxes within the oceanic water column fully. Recent advancements in remote sensing offer promising avenues for addressing these challenges. Studies investigating polarized radiance distribution and Chromophoric Dissolved Organic Matter (CDOM) provide valuable insights into improving remote sensing capabilities. Building upon these advancements, we propose a novel data-driven approach utilizing HyperOCR sensors and non-negative matrix factorization (NMF). Non-negative matrix factorization (NMF) is a powerful tool for extracting meaningful biological signatures from hyperspectral data, offering a granular yet comprehensive view of spectral diversity. Our study showcases the potential of NMF in elucidating spatial and temporal variations in biogeochemical processes within the ocean. Leveraging HyperOCR sensors, our approach offers a cost-effective and efficient means of enhancing remote sensing capabilities, enabling the rapid deployment and identification of seasonal patterns in the water column. Through extensive validation against field data from the Adriatic Sea, we demonstrate the utility of our approach in refining satellite measurements and improving algorithms for analyzing ocean color data. Our findings underscore the importance of integrating multiple observational platforms and advanced computational techniques to enhance the accuracy and reliability of remote sensing in ocean biogeochemistry studies. In conclusion, our study contributes to a deeper understanding of marine ecosystems’ responses to environmental changes and offers a new perspective on remote sensing capabilities, particularly in challenging coastal waters. By bridging the gap between satellite and in situ measurements, our approach exemplifies a promising pathway for advancing remote sensing of ocean biogeochemistry.

## INTRODUCTION

To understand the role of the ocean in climate change, it is important to interpret the biogeochemical fate of carbon in the ocean correctly. It has only recently become clear that the vast majority of the ocean carbon (662 Pg C) is found within the dissolved organic matter (DOM) pool, most of it in the reduced, refractory form [1]. Yet many questions about its role in climate feedback remain open, primarily regarding its potential remineralization by microbes or photo-oxidation (photoproduction of CO_2_)[2]. To fully understand it, spatially and temporarily, on a global scale, we need a combination of remotely obtained (satellite) and measured *in situ* relevant data through the entire water column [3].

Currently, the main needs cluster around developing and enhancing satellite radiation products to better support various research and operational applications related to ocean biology and biogeochemistry [4]. The ongoing challenges focus primarily on understanding complex oceanic processes and the increasing demand for precise and reliable data to inform environmental policy and management strategies, particularly in the face of climate change. The requirements include additional satellite-derived products such as sub-surface planar and scalar irradiance, average cosine, spectral fluxes (from UV to visible), diurnal fluxes, absorbed fraction of PAR by live algae (APAR), surface albedo, vertical attenuation, and heating rate. These products would provide more detailed and comprehensive data for studying marine ecosystems and their responses to environmental changes [5].

Despite the tremendous effort undertaken, we still lack information about subsurface processes governing carbon fluxes within the oceanic water column. The main problem remains that subsurface processes can only be detected remotely if they have a surface signature. With that in mind, various research approaches have been attempted to understand better and interpret satellite remote sensing capabilities.

Gleason et al. [6] detailed the measurement and modeling of the polarized upwelling radiance distribution in clear and coastal waters. The study successfully modeled and measured the Degree of Linear Polarization (DOLP) of the upwelling light field using a Monte Carlo-based radiative transfer code and fish-eye cameras equipped with linear polarizing filters. Field experiments in varying water conditions showed the model could predict the DOLP with an absolute error of ±0.05. This accuracy was achieved even with a fixed scattering Mueller matrix, which required precise *in situ* measurements of other optical properties [6]. The findings underscore the sensitivity of satellite sensors to polarization and the potential of using polarized radiance measurements for determining particle characteristics in oceanic waters [7,8]. The findings have significant implications for satellite remote sensing of the ocean floor by enhancing the accuracy and reliability of remote sensing data. Furthermore, Aurin et al. [9] demonstrate significant advancements in remote sensing of Chromophoric Dissolved Organic Matter (CDOM), CDOM spectral slope, and Dissolved Organic Carbon (DOC) in the global ocean. A comprehensive Global Ocean Carbon Algorithm Database (GOCAD) was developed using data from over 500 oceanographic field campaigns spanning three decades. This database incorporates a vast range of in situ reflectances, satellite imagery, and multispectral CDOM absorption coefficients, which facilitated the development, optimization, and validation of various semi-analytical, empirical, and machine learning algorithms for retrieving global DOC, CDOM, and CDOM slope. These algorithms have been optimized for global retrieval and exhibit a strong correlation with seasonal patterns of phytoplankton biomass and terrestrial runoff, highlighting their sensitivity and utility in understanding large-scale oceanic and atmospheric phenomena, such as the El Niño Southern Oscillation. Further validation of these algorithms, particularly in mid-ocean gyres and the Southern Oceans, is suggested to refine their application and increase accuracy. To address this problem, significant insights from a field intercomparison of radiometer measurements in the northern Adriatic Sea have been conducted to validate ocean color remote sensing data [10]. The study assessed the accuracy of in-water and above-water radiometer systems using multiple measurement systems under stable conditions. The results indicated generally good agreement among sensors for measuring downwelling irradiance, sky radiance, and above-water upwelling radiance, with differences typically less than 6% across visible wavelengths. The study further demonstrated the importance of accurate sensor calibration and highlighted the variability introduced by different measurement setups and environmental conditions. These findings are crucial for refining remote sensing methodologies, enhancing the reliability of ocean color data from satellite observations, and constraining the differences between in-water and satellite ocean color products. A similar study was conducted with the same goal, e.g. [11].

To tackle the disadvantages of remote sensing, comprehensive insights into the accuracy and challenges of *in situ* Ocean Color Radiometry (OCR) have been provided for the Southern Atlantic and Southeastern Pacific [12]. The research identified significant variability in remote sensing reflectance measurements obtained using different techniques, with relative percent differences ranging from 12% to 26% for ocean-color bands. This variability underscores the difficulties in obtaining precise *in situ* OCR measurements, particularly in regions with complex water properties and variable environmental conditions. The study also highlighted the critical impact of these uncertainties on the retrieval of *chlorophyll* concentrations and inherent optical properties using operational bio-optical algorithms. These findings have important implications for satellite remote sensing, particularly in calibrating and validating satellite sensors and improving bio-optical models for interpreting satellite data. The research emphasizes the need for refined measurement techniques and algorithms to enhance the accuracy of satellite-derived ocean color data, which is crucial for understanding global biogeochemical cycles and assessing climate change impacts on marine ecosystems. Finally, the advent of cutting-edge technology has paved the way for advancements in artificial intelligence and sensor systems. As these sensors become increasingly affordable, they are becoming more accessible to a wider range of applications. One such application is the aforementioned radiance in water columns, a task traditionally performed using satellite imagery. However, satellite data collection is fraught with challenges, including cloud cover, resolution limitations, orbit speed, and a lack of precision. Moreover, satellites fail to provide information about the entire water column, leaving a significant gap in our understanding.

To address these issues, we propose a data-driven approach that leverages HyperOCR sensors and artificial intelligence, specifically non-negative matrix factorization (NMF) (Figure 1A), as an alternative to previous approaches. NMF is a dimensionality reduction and data representation method that decomposes a given non-negative matrix into two lower-dimensional matrices, where all elements are constrained to be non-negative, enabling intuitive and interpretable parts-based representations of the original data [13](Figure 1B). NMF has been employed across various scientific fields and has demonstrated significant results in numerous studies [14–18]. Moreover, in recent studies on understanding the ocean’s carbon cycle, NMF-extracted signatures have shown an association between the abundance of microorganisms, providing further evidence of its potential utility [19]. Our approach is cost-effective and quick to deploy, offering a promising alternative to current methods revealing seasonal patterns in the water column (Figure 1C).

**Figure 1.**
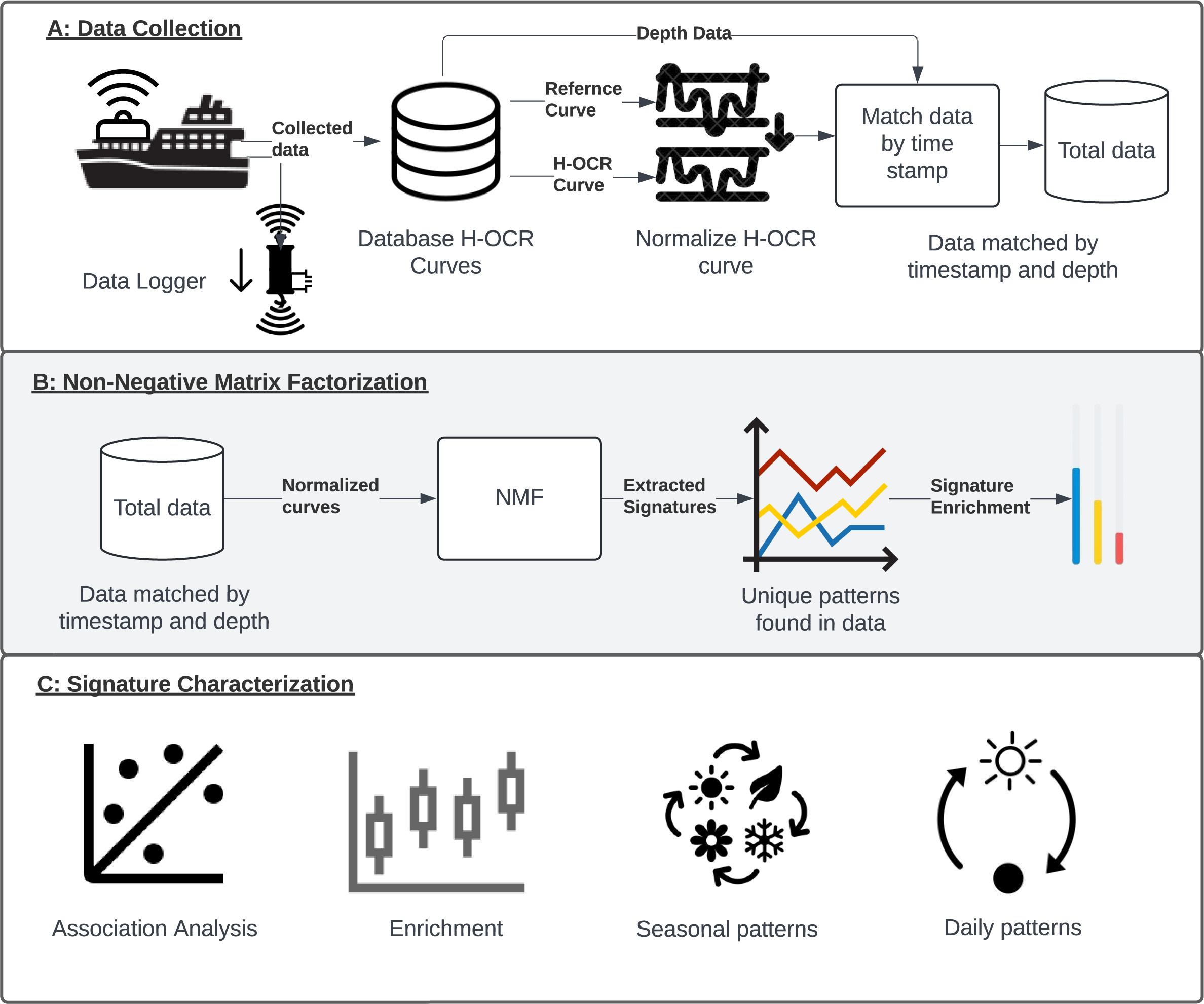
Study overview. A) Data collection process: Vessel equipped with sensor system was collecting data which consisted of H-OCR curves (UW and DW), temperature, pressure and reference curve(used for normalization). Finally the data was matched by the timestamp in order to have total dataset. B) NMF application: Normalized H-OCR curves were used in NMF with focus on extraction of unique patterns in data. C) Signature characterization: After signatures were disccovered by the NMF method, they were used in multiple downstream analyses.

## METHODS

### Data collection

We employed a custom-built profiler for our data collection, incorporating four sensors and a custom-built data logger (water-column profiler). Within this system, three sensors and a data logger were integrated into the profiler frame, while the fourth was positioned vertically on the vessel. Among the custom-built sensor system, two Hyperspectral Ocean Color Radiometers (Seabird HOCR) were deployed to capture both upwelling (UW) and downwelling (DW) irradiance. This arrangement facilitated irradiance measurement from both the upward and downward directions. Additionally, a Seabird SBE 39plus sensor was utilized to monitor pressure and temperature (Figure 1A). The fourth sensor, an Apogee PS-200 (Apogee Instruments, Inc., North Logan, UT, USA), served as a reference, capturing reference curves positioned atop the vessel. These sensors provided comprehensive data acquisition capabilities, enabling detailed monitoring and analysis throughout our study.

### Experiment location and sampling

The experiment was conducted near the islands of Mljet (42L 45′ 15″ N; 17L 23′ 12″ E, depth 128 m) and Vis (43L 03’ 32.8’’ N, 16L 17’ 19.7’’ E, depth 102.9 m) in Croatia. All measurements were taken in the middle of the day, at the sun’s culmination. The profiler’s descending speed was 0.22 m/s. A total of 22 profiles were analyzed, resulting in the collection of 11,653 hyperspectral curves (Figure 2A). These measurements were conducted intermittently across May, June, July, August, and December to capture data potentially influenced by seasonal patterns (Figure 2B).

**Figure 2.**
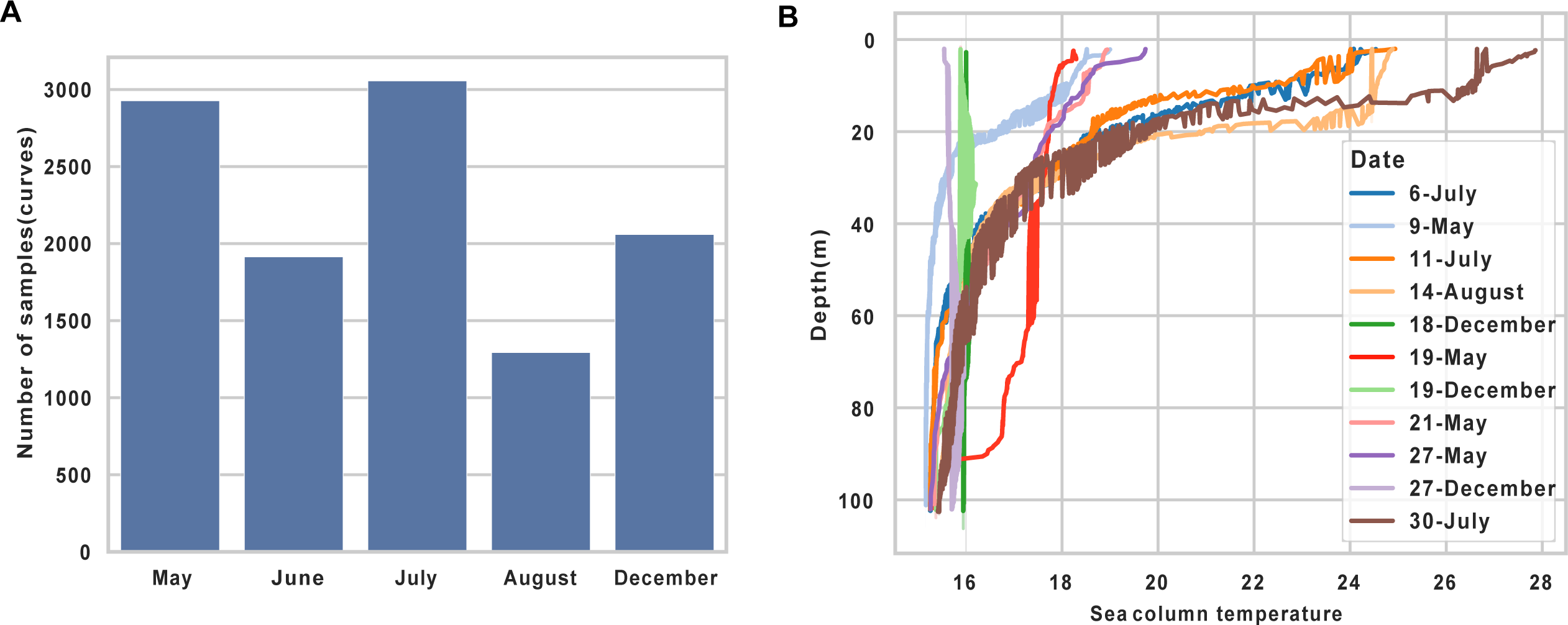
Description of data. A) Barplot shows number of H-OCR curves collected per month. B) Line plot shows association between depth (y-axis) and water column temperature in Celsius (x-axis) for every day in this study. Red line indicating the measurement conducted on 19th of May shows consistent temperature across almost entire water column.

### Data preprocessing and normalization

The data underwent preprocessing and normalization through the following steps. Initially, all sensor data were merged based on timestamps, allowing a tolerance of up to 2 seconds to accommodate variations in sampling rates among sensors. Next, the Apogee sensor served as a reference and was utilized for normalization purposes. Hyperspectral curves from the upwelling (UW) and downwelling (DW) sensors were normalized by dividing them by the reference curve (Figure 1A). Furthermore, the dataset was filtered to include only curves measured at a depth of 1 meter or greater, as sensors require time to auto-calibrate when submerged in the water column.

### Application of Non-Negative Matrix Factorization

The Non-Negative Matrix Factorization (NMF) algorithm decomposes an input matrix into two non-negative matrices, W and H, such that their product approximates the input matrix (Supplementary Figure 1). The matrix W represents the basis or dictionary elements, often interpreted as unique patterns or features present in the data, such as hyperspectral ocean color radiometer (h-OCR) curves, which we refer to as “signatures”. The matrix H represents the coefficients or weights that determine the contribution of each basis element to reconstruct the original data curves. In our implementation of the NMF algorithm, we utilized the ‘nndsvdar’ initialization method to compute the initial state of the factorization. This initialization approach is based on the Non-negative Double Singular Value Decomposition with Alternating Rectification (NNDSVDar) method, which initializes the factor matrices with small random values, allowing for faster convergence. The NMF solver employed multiplicative updates iteratively to optimize the factor matrices while minimizing the Frobenius norm as the loss function. We set the maximum number of iterations for the NMF algorithm to 1000 to ensure convergence to a satisfactory solution. This comprehensive approach enabled us to effectively decompose the input matrix into meaningful basis patterns and their corresponding coefficients, facilitating the extraction of interpretable features from the data.

### Code Availability

The code for analysis and plotting is accessible via the public GitHub repository at https://github.com/mxs3203/svjetloPaper. The repository contains code for the NMF model, HOCR data processing, and figure generation, which are included in the manuscript. Additionally, the data matrices are available for download as Supplementary Material.

### Computational requirements

Computational modeling was performed using a standard desktop computer with average performance. As the modeling process did not utilize a GPU and did not require extensive computational resources, a regular computer sufficed for our purposes.

## RESULTS

### Temperature measurements show patterns of water column stratification

The temperature-depth profile depicted in Figure 2B illustrates temperature distribution across the water column during sampling intervals (Supplementary Figure 2). Results indicate gradual stratification of the water column from May to August. In December, the water column was vertically homogeneous with a temperature of 16°C as a consequence of wind Bora mixing. Measurements conducted on the 19th of May 2023 (Figure 2B, Supplementary Figure 3) show an inflow of Ionian Surface Water (ISW) via the Otranto Strait to the depth of 95m, which is characteristic for autumn [20]. However, circulation in the Adriatic Sea exhibits high spatial and temporal variability, with a general cyclonic circulation pattern [21].

### Fitting the NMF model to hyperspectral data

In this study, we employed Non-Negative Matrix Factorization (NMF) to uncover underlying patterns within the normalized hyper-OCR curves dataset. The NMF algorithm decomposes the input data matrix into two non-negative matrices: a basis matrix (or signature matrix) and a coefficient matrix (Supplementary Figure 1). The basis matrix represents distinct signatures or patterns inherent in the data, while the coefficient matrix indicates the contribution of each signature to the original samples. Through the training process, our NMF model generated a set of signatures, each represented as a unique hyperspectral curve, encapsulating unique patterns within the dataset. These signatures serve as interpretable representations of the underlying structure of the data. To determine the optimal number of signatures (k) for our model, we employed the “elbow method”, a common approach for selecting the appropriate number of clusters or components in unsupervised learning tasks. Our analysis revealed that k=5 emerged as the optimal number of signatures (Figure 3A). Beyond this value, the reconstruction error did not exhibit substantial changes, suggesting that additional signatures did not capture significantly more variance in the data. Thus, we proceeded with k=5 to extract the most salient patterns from the hyper-OCR curves dataset. NMF analysis yielded five distinct signatures (S1 to S5), each representing unique spectral patterns within the dataset(Figure 3B). Signature 1 (S1) is characterized by a high-intensity peak centered around 440nm. Signature 2 (S2) exhibits a high-intensity peak starting at 350 nm and rapidly decreasing towards 450nm, with an additional small peak between 500 nm and 600 nm, reaching its highest intensity at 580 nm. Signature 3 (S3) presents a broad spectral curve with medium intensity spanning from 500 nm to 700 nm, with its peak intensity observed at approximately 585 nm. Signature 4 (S4) is defined by a medium-intensity peak around 460nm and shows a small enrichment in the spectral range of 800 nm to 850nm. Finally, Signature 5 (S5) displays a medium-intensity peak resembling a bimodal function, with peaks at 495 nm and 520nm (Figure 3B).

**Figure 3.**
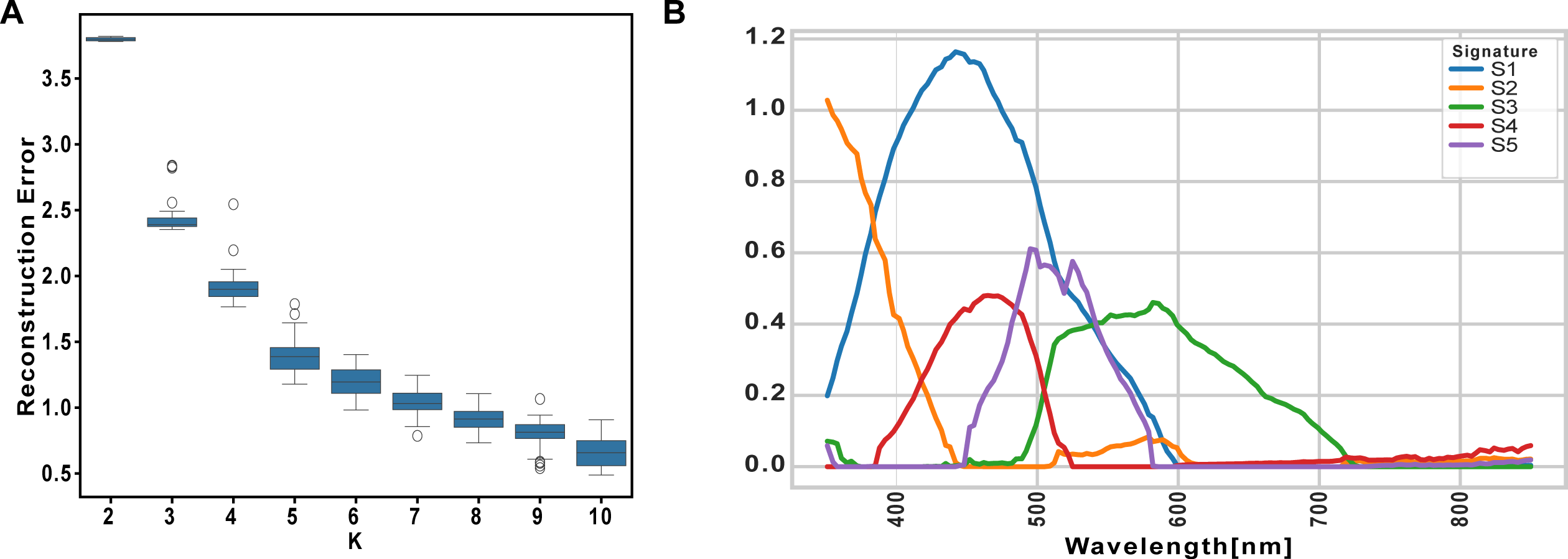
NMF Application. A) Reconstruction error for each number of signatures (K). Boxplot indicates that error decreases rapidly until reading K=5, indicating optimal K. B) Visualization of extracted signatures.

### Comparing UW and DW sensor signatures across months

We divided the signatures obtained from the upwelling (UW) and downwelling (DW) sensors for separate analyses. The UW sensor exhibited the lowest values in signatures S1, S3, and S5, while the DW sensors demonstrated the highest values in S1 and S4 (Figure 4A-B). Upon examining the UW sensor signatures from May to December, we observed a gradual increase in S2, S3, and S4, with peaks occurring in July and August before returning to lower values in December. Notably, S1 started with the highest values in May and gradually decreased towards December. Additionally, S5 showed minor enrichment in the UW sensor, with small peaks observed in May and December (Figure 4A). The downwelling (DW) sensor generally exhibited the highest enrichment values in S1, with peaks observed in May and August. S2 demonstrated comparable values in May and June, gradually increasing towards August and December. S3 exhibited the highest values in December and May, displaying a U-shaped pattern with a minimum observed in July and August. High enrichment in S4 was consistently observed in the DW sensor, peaking in June, July, and August. Lastly, S5 displayed comparable enrichment levels in May, June, and July, with a noticeable increase observed in August and December (Figure 4B).

**Figure 4.**
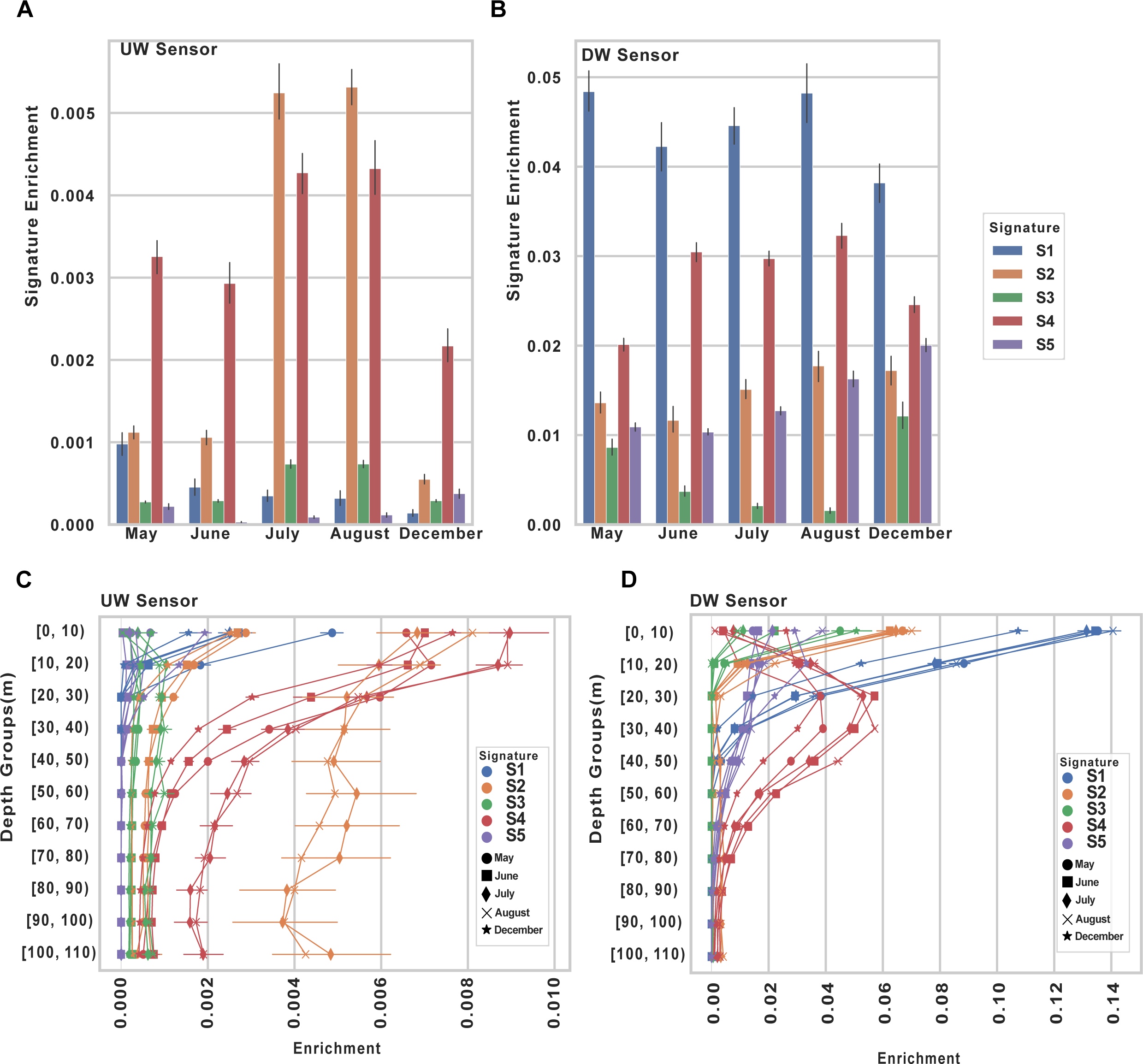
Signature characterization. A) Barplot indicating signature enrichment for UW sensors for each month. B) Bar plot indicating signature enrichment for DW sensor for each month. C) Line plot indicating average enrichment of signatures from UW sensor for specific depth bin and month. D) Line plot indicating average enrichment of signatures from DW sensor for specific depth bin and month.

### Exploring depth and monthly variations in sensor signatures

In this analysis, we categorized depth measurements into bins spanning intervals of 10 meters each, starting from 0 to 10 meters and extending onwards in increments of 10 meters. Our objective was to identify any trends in signature enrichment across different depths and months. In the upwelling (UW) sensor data, we observed consistent patterns across most months for signature S1, except in May, where it exhibited the highest average enrichment (Figure 4C, Supplementary Figure 4). However, this enrichment decreased as depth increased. For S2, similar patterns were noted in July and August, with noticeable differences between depths of 50 meters and 90 meters, where July showed higher enrichment (Figure 4C). Notably, S2 did not exhibit the same rapid decrease with depth in July and August as observed in other months as signature enrichment stayed stable. Conversely, S3 and S5 displayed minimal enrichment and exhibited minor variations across the study months. Nonetheless, it is noteworthy that S5 displayed the highest enrichment in December within depths of 0 to 10 meters, gradually decreasing with increasing depth. In S4, comparable patterns were observed in July and August, while May, June, and December displayed distinct decreasing patterns. December exhibited the steepest decrease in S4 enrichment. In the downwelling (DW) sensor data, distinct patterns emerged, particularly in the downward trend of signature enrichment with increasing depth, observed across all signatures except S4 (Figure 4D, Supplementary Figure 4). S1 exhibited a consistent downward trend in enrichment across all months except for December, when the downward pattern was a bit steeper. S3 displayed comparable trends in December-May and July-August, with June standing out as having a distinct pattern. The pattern observed in S4 is particularly interesting, where enrichment initially began with values close to zero near the surface, except for December. Subsequently, the enrichment of S4 gradually increased, peaking between 30 and 40 meters depth before declining as depth increased(Figure 4D). S5 demonstrated an almost linear downward pattern, starting with the highest enrichment near the surface and gradually decreasing with depth, with notable differences observed in December and August. In December, S5 exhibited higher values around 10 and 20 meters than the average values at 0 to 10 meters.

### Daily patterns of sensor enrichment

To assess daily patterns, we categorized sampling time into three distinct periods. “Morning” was defined as the time frame between 6:00 AM and 11:00 AM, “Noon” encompassed the period from 11:00 AM to 5:00 PM, and “Dusk” extended from 5:00 PM to 9:00 PM. These categories were employed to group sensor enrichment data and calculate the average enrichment of signatures across each designated time period for both sensors. In the UW sensor data, signatures S1, S3, and S5 exhibited minimal changes across the three defined time frames and months. However, S2 displayed distinct patterns, particularly in August, indicating that enrichment was highest in the morning and lowest at dusk. In May, S2 showed an upward pattern, with increased values observed at dusk. Notably, in July, S2 displayed the lowest values at noon but experienced a drastic increase in signature enrichment at dusk. Similarly, S4 demonstrated similar patterns in July, with the highest enrichment observed at dusk. In June, S4 exhibited identical enrichment levels in the morning and noon but showed zero enrichment at dusk, while December showed the highest enrichment of S4 at noon (Figure 5A). In the DW sensor data, distinct patterns emerged, with notable differences observed between the months and time frames used for analysis. S1 exhibited the highest enrichment at dusk in May and July, while S2 showed the highest enrichment in the morning in August and July. Interestingly, S2 in July also displayed the highest enrichment at dusk. S3 did not demonstrate any significant patterns or major changes across months or time frames. On the other hand, S4 exhibited the highest and similar enrichment levels in the morning and noon in June, July, and August, with differences observed at dusk. In July, S4 maintained high enrichment levels at dusk, whereas in June and July, enrichment levels diminished to almost zero. Additionally, S4 displayed notable results for December, with the highest enrichment observed at noon. Similarly, S5 also showed the highest average enrichment at noon in December. August and June exhibited similar patterns, with the highest enrichment observed in the morning, gradually decreasing towards dusk (Figure 5B).

**Figure 5.**
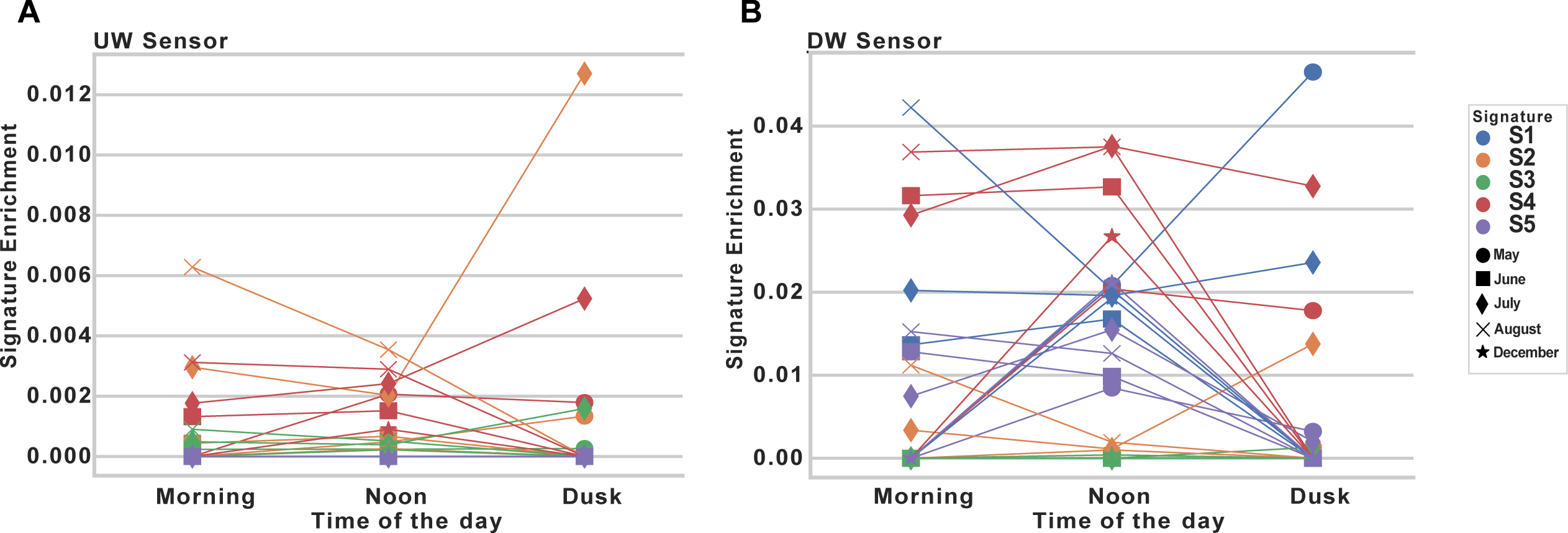
Signature enrichment by time intervals. A) Line plot indicating how enrichment of signatures in UW sensor, changes during the time of day and month. B) Line plot indicating how enrichment of signatures in DW sensor, changes during the time of day and month.

## DISCUSSION

Our study utilized a custom-built sensor system to collect comprehensive data, resulting in a dataset comprising 11,653 hyperspectral curves paired with temperature and depth measurements. While conducting the sampling and comparing the data afterward, we observed differences in the quality of measurements during specific time frames and depth ranges. Particularly, measurements taken in the late afternoon, approximately 1-2 hours before dusk, exhibited a notable amount of noise. Additionally, we observed the influence of wave activity on the quality of measurements, particularly in the surface layer. Waves induced significant noise in the measurements due to light scatter, resulting in inconsistencies and inaccuracies (Supplementary Figure 5). However, we found that as depth increased, typically between 10-20 meters, the influence of wave-induced noise decreased, leading to more stable and reliable measurements. Both problems were addressed at the data filtering step, where noisy data was removed. The temperature data revealed intriguing patterns, particularly observed on the 19th of May, 2023, wherein a homogeneous temperature distribution was observed with decreasing depth. This event relates to the inflow of Ionian Surface Water (ISW) via the Otranto Strait to a depth of 95m, which is normally characteristic of autumn [20].

Employing a Non-Negative Matrix Factorization (NMF) model on the entire dataset yielded five distinct signatures. These signatures were instrumental in characterizing both seasonal and daily patterns within our data. The differentiation in spectral signatures between upwelling (UW) and downwelling (DW) sensors, especially their distinct seasonal dynamics, can be partially explained by the variations in phytoplankton community composition and its biogeochemical implications. For instance, the highest enrichment in signature S1 during the peak of light upwelling values and its gradual decrease could be indicative of phytoplankton dynamics where certain taxa like diatoms dominate due to nutrient influxes, consistent with findings that growth conditions can decouple diatom populations from grazing pressure, leading to significant bloom events [22]. This hypothesis is supported by our observation of spectral signature shifts that coincide with known nutrient dynamics and light conditions, which are critical drivers of phytoplankton community structure [22,23]. Interestingly, the variation in signatures across different depths and months, particularly the observed depth-wise attenuation of signature enrichment in DW sensor data, underscores the stratification effects on light availability and nutrient distribution, aligning with observations that certain cyanobacteria and smaller phytoplankton are adapted to low-nutrient, high-light conditions in stratified waters [24]. The consistent patterns seen in certain signatures during specific months could also reflect the physiological adaptations of phytoplankton to seasonal light variations, potentially impacting their backscattering properties as suggested by the taxonomic variability in particle backscattering (BBP)-to-phytoplankton carbon (Cphyto) scaling [25]. Moreover, our findings add a nuanced layer to understanding phytoplankton dynamics by linking spectral data with ecological patterns. For example, the enrichment peaks in signatures from the UW sensor during early summer might indicate rapid growth phases or succession events within the phytoplankton community, which are critical for predicting carbon export potential during these periods. This aligns with the discourse on the impact of community composition on carbon cycling and the necessity of considering taxonomic variability in modeling biogeochemical cycles [22,26]. Future work should focus on integrating these spectral signatures with direct taxonomic identification and physiological state assessments to validate the inferred patterns and enhance the predictive power of hyperspectral analyses in marine ecosystems. Additionally, adopting newer, absorption-based methods for estimating phytoplankton biomass and carbon content could refine the interpretation of spectral data, especially in relation to community composition changes and their biogeochemical roles [27]. Our study underscores the variability and complexity of phytoplankton dynamics in marine ecosystems and highlights the potential of sophisticated analytical techniques like NMF in unraveling these complexities. By advancing our understanding of how different phytoplankton communities contribute to and are influenced by environmental factors, we can better predict and manage the impacts of global changes on marine ecosystems. As global oceanic and atmospheric conditions continue to evolve, significant shifts in phytoplankton community structure are anticipated, particularly with respect to picophytoplankton and cyanobacteria. These groups, characterized by their small size and adaptability, are poised to dominate increasingly stratified and oligotrophic waters driven by rising sea surface temperatures and reduced nutrient inputs from deep waters [23,28]. Cyanobacteria, in particular, are known for their efficiency in low-nutrient environments and could outcompete larger phytoplankton taxa under such conditions, potentially leading to a higher prevalence of cyanobacteria in future marine ecosystems [29]. This shift could have profound implications for marine food webs and biogeochemical cycles, as cyanobacteria and other picophytoplankton typically have different nutritional contents, sinking rates, and interactions with higher trophic levels compared to larger phytoplankton like diatoms. Moreover, the increase in cyanobacteria may also influence the biogeochemical properties of marine environments, such as nitrogen fixation rates and carbon sequestration capabilities. Predictive models need to account for these changes to accurately forecast the impacts of climate change on marine ecosystems, particularly the potential feedback mechanisms involving picophytoplankton that could alter oceanic carbon cycling dynamics and nutrient fluxes significantly. We believe that our approach to studying light in the water column, as presented in this study, could significantly add to a better understanding of these changes.

## FUNDING

This research was funded by Regionale Forskningsfond Agder (RFF Agder), project number 338390. The development of the new technology used in this study was funded by Innovasjon Norge (Innovation Norway).

## ACKNOWLEDGEMENTS

We would like to thank the Ministry of Environmental Protection and Energetics (Ministarstvo zaštite okoliša i energetike) of the Republic of Croatia and Mljet National Park, Croatia for their support and for making this study possible; we also thank Adrián Gómez Repollés for his comments on the manuscript. We would also like to thank Ivana and Anka Hajdić for the use of their boat and help with data collection.

---

**Supplementary** Figure 1: Visalization of NMF factorization process for using our data

**Supplementary** Figure 2: Distribution of temperature measurements for each month

**Supplementary** Figure 3: Line plot showing temperature distribution across water column (depth).

**Supplementary** Figure 4: Barplots showing average enrichment of signatures per sensor, month and depth-bin

**Supplementary** Figure 5: Example of tidy signature extraction due stable sea(left) vs. signature extraction when light was scattered due waves (right)

## Notes

### Competing Interest Statement

The authors have declared no competing interest.

