## Supplementary figures and images for "Advancing Oceanic Studies with HyperOCR Sensors and Non-Negative Matrix Factorization: A Cost-Effective, Data-Driven Approach for Analyzing Light in Marine Water Column"

### Supplementary Figure 1

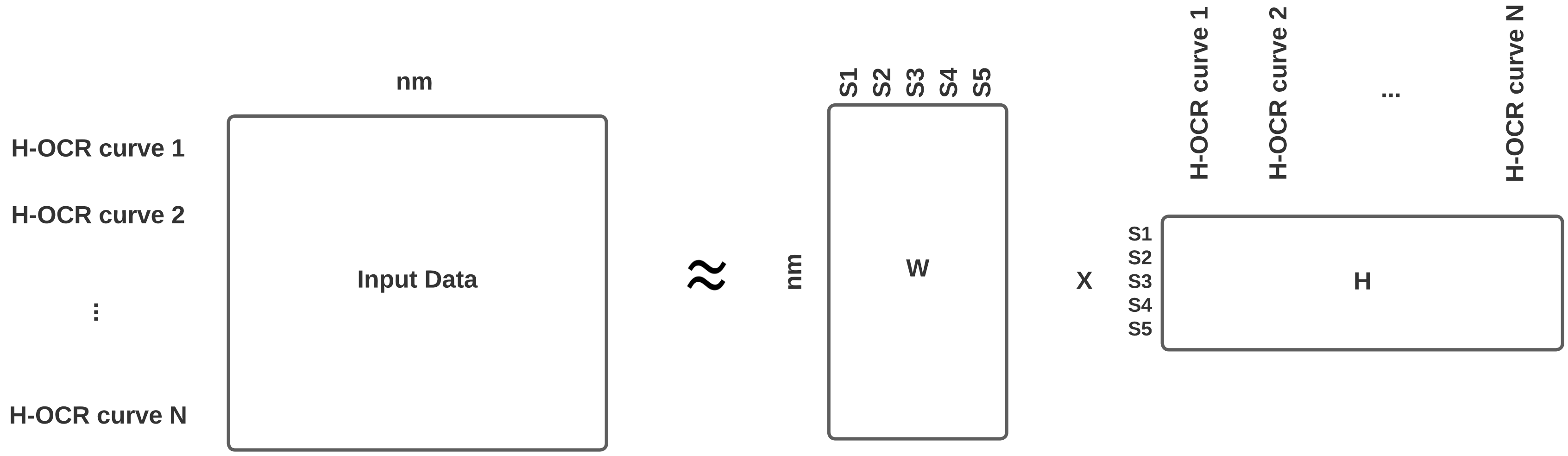

### Supplementary Figure 2

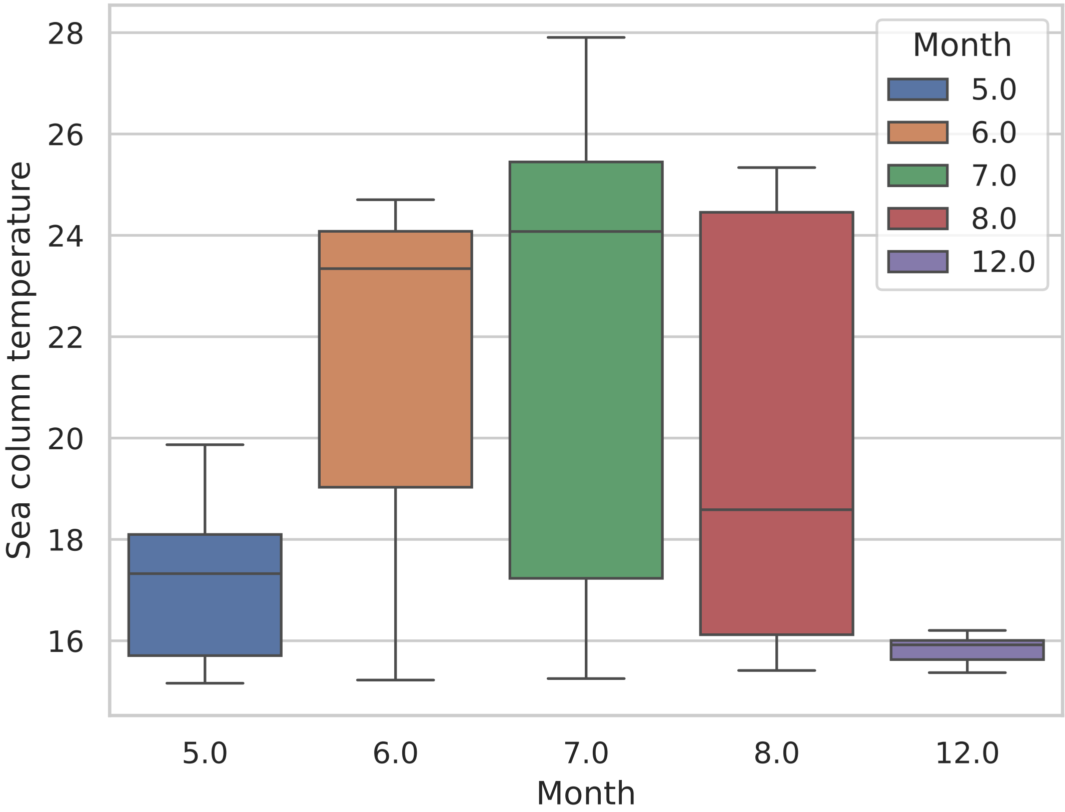

### Supplementary Figure 3

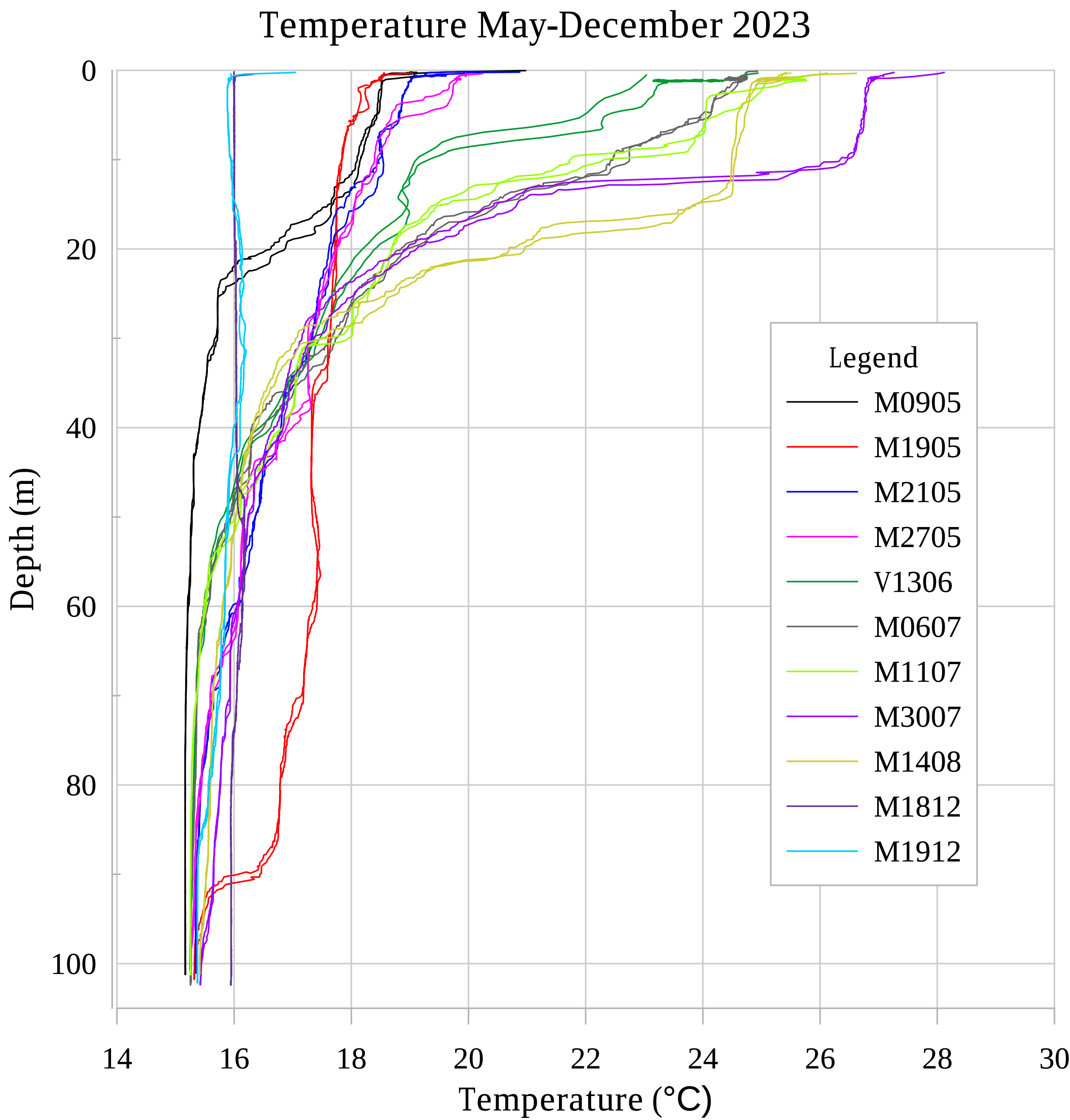

### Supplementary Figure 4

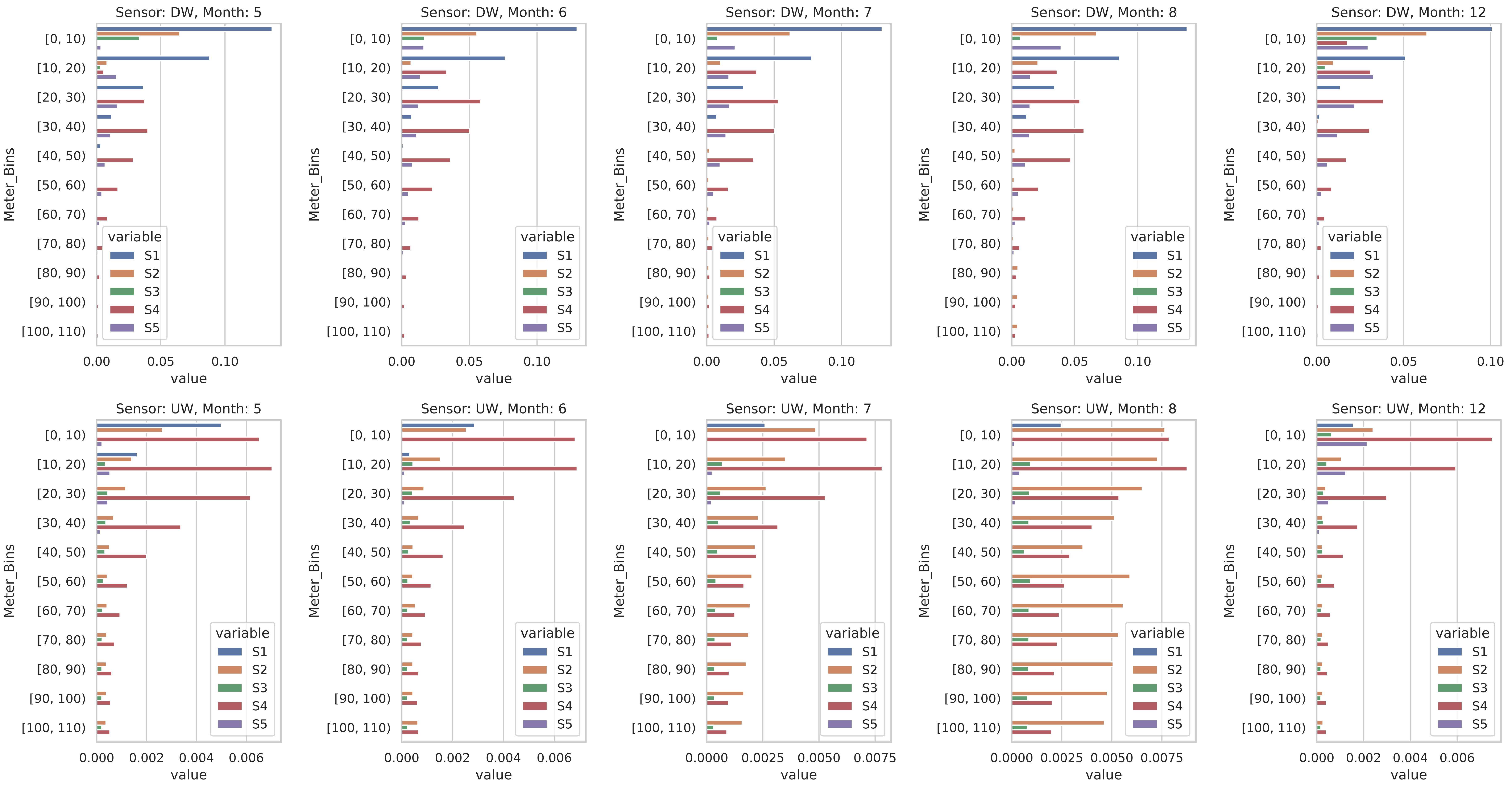

### Supplementary Figure 5

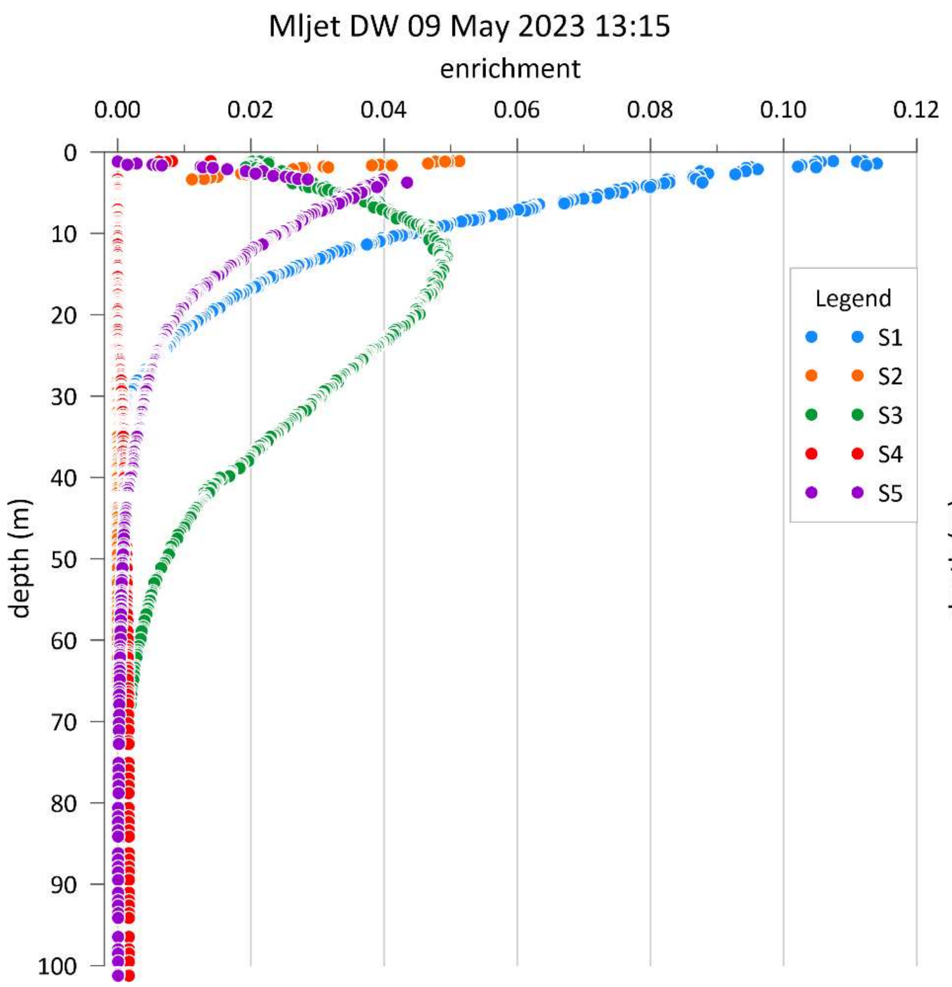

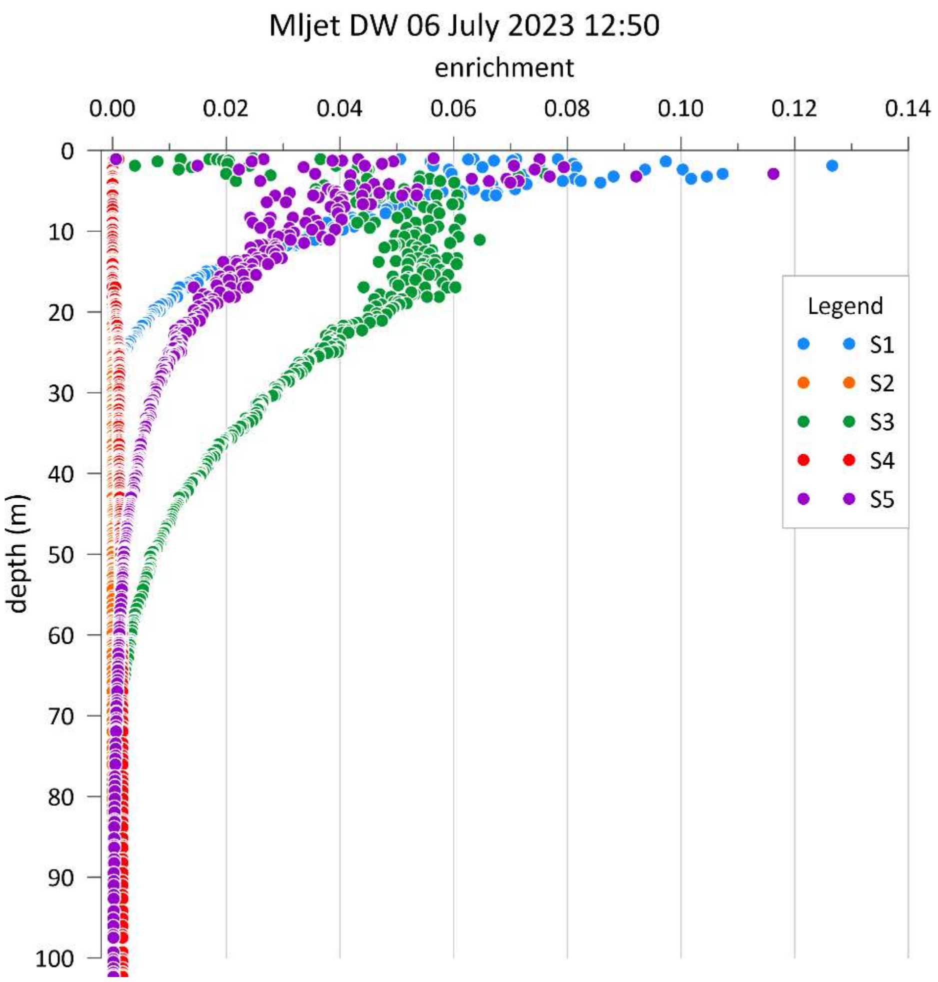
